# Nested Stochastic Block Models Applied to the Analysis of Single Cell Data

**DOI:** 10.1101/2020.06.28.176180

**Authors:** Leonardo Morelli, Valentina Giansanti, Davide Cittaro

## Abstract

Single cell profiling has been proven to be a powerful tool in molecular biology to understand the complex behaviours of heterogeneous system. The definition of the properties of single cells is the primary endpoint of such analysis, cells are typically clustered to underpin the common determinants that can be used to describe functional properties of the cell mixture under investigation. Several approaches have been proposed to identify cell clusters; while this is matter of active research, one popular approach is based on community detection in neighbourhood graphs by optimisation of modularity. In this paper we propose an alternative and principled solution to this problem, based on Stochastic Block Models. We show that such approach not only is suitable for identification of cell groups, it also provides a solid framework to perform other relevant tasks in single cell analysis, such as label transfer. To encourage the use of Stochastic Block Models, we developed a python library, `schist`, that is compatible with the popular `scanpy` framework.

## Background

Transcriptome analysis at single cell level by RNA sequencing (scRNA-seq) is a technology growing in popularity and applications [1]. It has been applied to study the biology of complex tissues [2, 3], tumor dynamics [4, 5, 6, 7], development [8, 9] and to describe whole organisms [10, 11].

A key step in the analysis of scRNA-seq data and, more in general, of single cell data, is the identification of cell populations, that is groups of cells sharing similar properties. Several approaches have been proposed to achieve this task, based on well established clustering techniques [12, 13], consensus clustering [14, 15, 16] and deep learning [17]; many more have been recently reviewed [18, 19] and benchmarked [20]. As the popularity of single cell analysis frameworks `Seurat` [21] and `scanpy` [22] raised, methods based instead on graph partitioning became the *de facto* standards. Such methods require the construction of a cell neighbourhood graph (*e.g.* by *k* Nearest Neighbours, *k*NN, or shared Nearest Neighbours, *s*NN). Encoding cell-to-cell similarities into graphs has practical advantages beyond clustering, as many algorithms for graph analysis can be applied and interpreted in a biological way. A notable example is the analysis of cell trajectories which can be derived from the analysis of Markov processes traversing the NN graph [23, 24]. In another context, computation of RNA moments in scRNA velocity is also based on the NN graph structure [25]. Arguably, the biggest utility of NN structure is the possibility to identify cell groups by partitioning the graph into communities; this is typically achieved using the Louvain method [26], a fast algorithm for optimisation of graph modularity. While fast, this method does not guarantee the identification of internally connected communities. To overcome its limits, a more recent approach, the Leiden algorithm [27], has been implemented and it has been quickly adopted in the analysis of single cell data, for example by `scanpy` [22] and `PhenoGraph` [28]. In addition to Newman’s modularity [29], other definitions currently used in single cell analysis make use of a resolution parameter [30, 31]. In lay terms, resolution works as a threshold on the density within communities: lowering the resolution results in less and sparser communities and *vice versa*. Identification of an appropriate resolution has been recognised as a major issue [32], also because it requires the definition of a mathematical property (clusters) over biological entities (the cell groups), with little formal description of the latter. In addition, the larger the dataset, the harder is to identify small cell groups, as a consequence of the well-known resolution limit [33]. Moreover, it has been demonstrated that random networks can have modularity [34] and its optimisation is incapable of separating actual structure from those arising simply of statistical fluctuations of the null model. Lastly, it is a common error to assume that the resolution parameter reflects a hierarchical structure of the communities in the graph when, in general, this is not rigorously true. Additional solutions to cell group identification from NN graphs have been proposed, introducing resampling techniques [35, 36] or clique analysis [37]. It has been proposed that high resolution clustering, *e.g.* obtained with Leiden or Louvain methods, can be refined in agglomerative way using machine learning techniques [38].

An alternative solution to community detection is the Stochastic Block Model, a generative model for graphs organised into communities [39]. In this scenario, identification of cell groups requires the estimation of the proper parameters underlying the observed NN graph. According to the microcanonical formulation [40], the parameters are partitions and the matrix of edge counts between them. Under this model, nodes belonging to the same group have the same probability to be connected together. It is possible to include node degree among the model parameters [41], to account for heterogeneity of degree distribution of real-world graphs. A Bayesian approach to infer parameters has been developed [42] and implemented in the `graph-tool` python library (https://graph-tool.skewed.de). There, a generative model of network ***A*** has a probability *P*(***A***|***θ***, ***b***) where ***θ*** is the set of parameters and ***b*** is the set of partitions. The likelihood of the network being generated by a given partition can be measured by the posterior probability

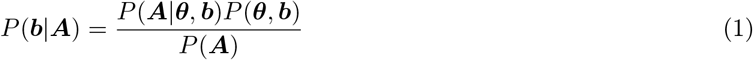

and inference is performed by maximising the posterior probability. The numerator in Eq. 1 can be rewritten exponentiating the description length

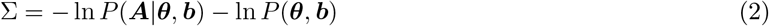

so that inference is performed by minimising the information required to describe the data (Occam’s razor); `graph-tool` is able to efficiently do this by a Markov Chain Monte Carlo approach [43]. SBM itself may fail to identify small groups in large graphs, hence hierarchical formulation has been proposed [44]. Under this model, communities are agglomerated at a higher level in a block multigraph, also modelled using SBM. This process is repeated recursively until a graph with a single block is reached, creating a Nested Stochastic Block Model (nSBM).

In this work we propose nSBM for the analysis of single cell data, in particular scRNA-seq data. This approach identifies cell groups in a statistical robust way and, moreover, it is able to determine the likelihood of the grouping, thus allowing model selection. In addition, it is possible to measure the confidence of assignment to groups, a measure that can be exploited in various analysis tasks.

We developed `schist` (https://github.com/dawe/schist), a python library compatible with `scanpy`, to facilitate the adoption of Stochastic Block Models in single-cell analysis.

## Results

### Overview of schist

`schist` is a convenient wrapper to the `graph-tool` python library, written in python and designed to be used with `scanpy`. The most prominent function is `schist.inference.nested_model()` which takes a `AnnData` object as input and fits a nested Stochastic Block Model on the *k*NN graph built with `scanpy` functions (e.g. `scanpy.tools.neighbors()`). When launched with default parameters, `schist` fits a model which maximises the posterior probability of having a set of cell groups (or blocks) given a graph. `schist` then annotates cells in the data object with all the groups found at each level of a hierarchy. Given the large size of the NN graph in real-world experiments, it is possible that a single solution represents local minima of the fitting process. In addition, it is possible that multiple solutions are equally acceptable to represent the graph partitioning and a better description is given by the consensus over such solutions [45]. To overcome these issues, `schist` fits multiple instances in parallel and returns the inferred consensus model, alongside the marginal probabilities for each cell to belong to a specific group (*cell marginals*). Moreover, the Stochastic Block Model has no constraints on what type of modular structure is fitted, meaning that groups are not necessarily identified only by assortativity (i.e. cells are mostly connected within the same group). When assortativity is thought to be the dominant pattern another model (the Planted Partition Block Model, PPBM [46]), also implemented in `schist`, is better suited to find statistically significant assortative communities, also eliminating the need to set a resolution parameter as required in standard community detection by maximisation of modularity.

### Absence of information is correctly handled by nSBM

One of the most relevant difference between SBM and other methods to cluster single cells is that it relies on robust statistical modelling. In this sense, the number of groups identified strictly mirrors the amount of information contained in the data. An important consequence is that absence of information can be properly handled. To show this property we performed a simple experiment on a randomised *k*NN graph. We collected data for 3k PBMC (available as preprocessed data in `scanpy`) and shuffled the edges of the prebuilt *k*NN graph, this to keep the general graph properties unchanged. We tested that the degree distribution does not change after randomisation (Kolmogorov-Smirnov *D* = 0.0733, *p* = 0.703). We found that the default strategy, based on maximisation of modularity, identifies 24 cell groups, whereas nSBM does not identify any cell group.

This experiment is a deliberate extreme case. The quality of grouping proposed by a standard approach can be disputed in many ways, and the UMAP embedding indeed reflects the absence of any information. Nevertheless, real-world data may include an unknown amount of random noise. Hence, It is important to identify cell groups that are not artefacts arising from processing and that do reflect the information contained in the dataset.

### schist correctly identifies cell populations

To benchmark `schist`, we tested it on scRNA-seq mixology data [47], a dataset explicitly developed to benchmark single cell analysis tools without the need to simulate data. In particular, we used the mixture of 5 cell lines profiled with Chromium 10x platform. At a first evaluation of the UMAP embedding, all lines appear well separated. Only the lung cancer line H1975 shows a considerable degree of heterogeneity and appears to be split into two cell groups (Fig. 2A). Using default parameters, `schist` is able to identify correct cell groups (*ARI* = 0.829), with a further split in H2228 cell line (Fig. 2B), whereas Leiden method clusters the dataset into 10 groups (*ARI* = 0.549, Fig. 2C). The nSBM solution correctly identifies H1975 groups as a single entity at level 1 of the hierarchy. We then sought to check if an independent agglomerative method, SCCAF [38], was able to recover cell line groupings starting from both partition schemes. Given the ground truth, the cell lines, SCCAF is able to assess the maximal accuracy that can be achieved in the dataset (0.992). When trained with this target accuracy, SCCAF precisely reconstructs the original cell line annotations starting from `schist` partitions with high accuracy (Fig. 2D). When Leiden partitions are set as input, SCCAF merges H2228 and HCC827 cells into a single cluster and keeps H1975 cells split into two groups (Fig. 2E), highlighting potential limitations of this approach. In order to perform a fair and complete experiment, we also tested SCCAF with the partitions scheme at the deepest level of the nSBM hierarchy (38 groups, Fig. S1A), for which we observed the inability to perform a significant cell grouping (Fig. S1B).

**Figure 1:**
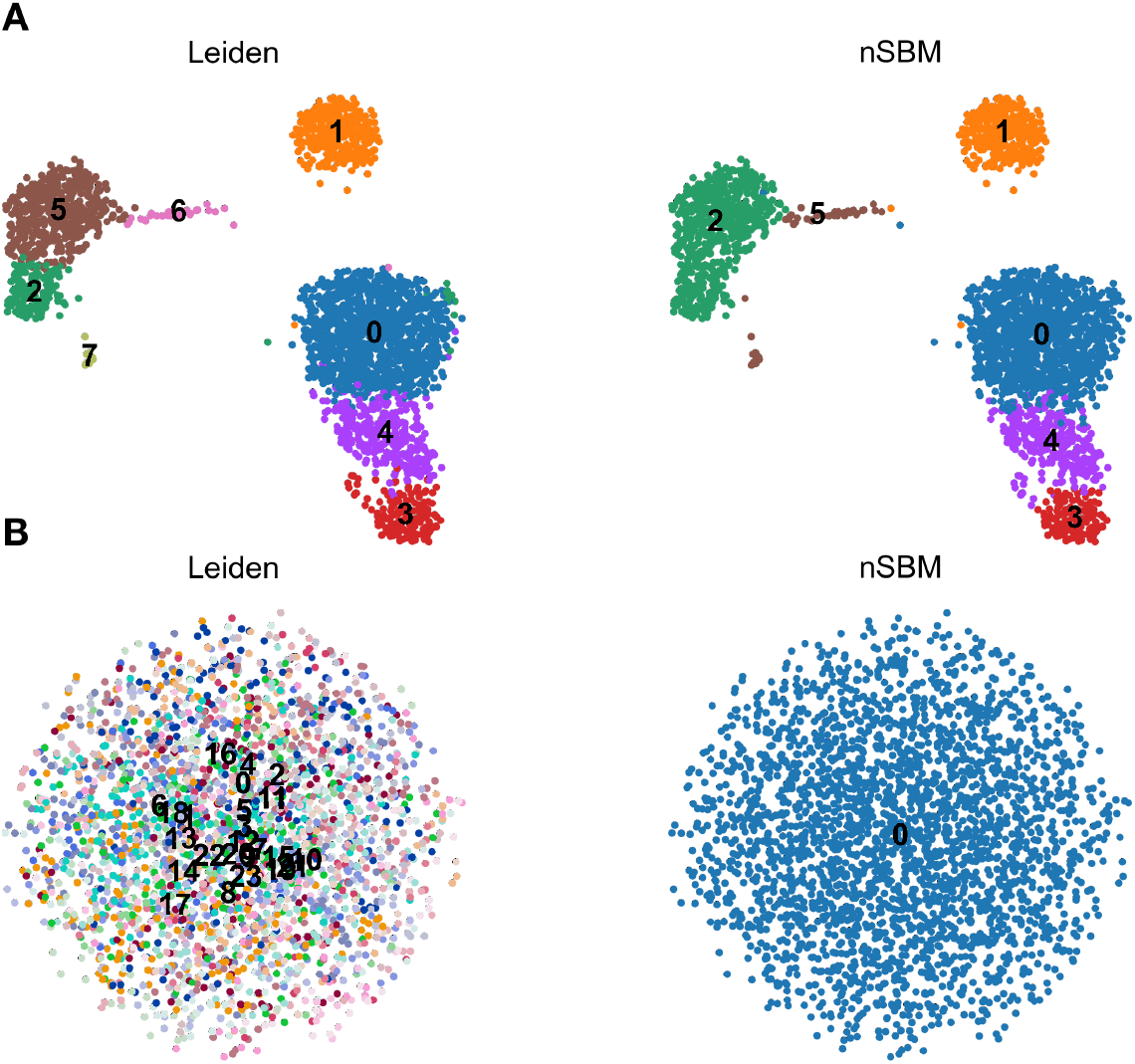
nSBM results for randomised data. (A) UMAP embedding of 3k PBMC after standard processing with Leiden approach (left) or nSBM (right). Cell grouping is consistent for both the approaches (Adjusted Rand Index *ARI* = 0.869). (B) UMAP embedding of the same data after randomisation of *k*NN graph edges. In this case nSBM does not return any cell grouping, while optimisation of modularity finds up to 24 different cell groups

**Figure 2:**
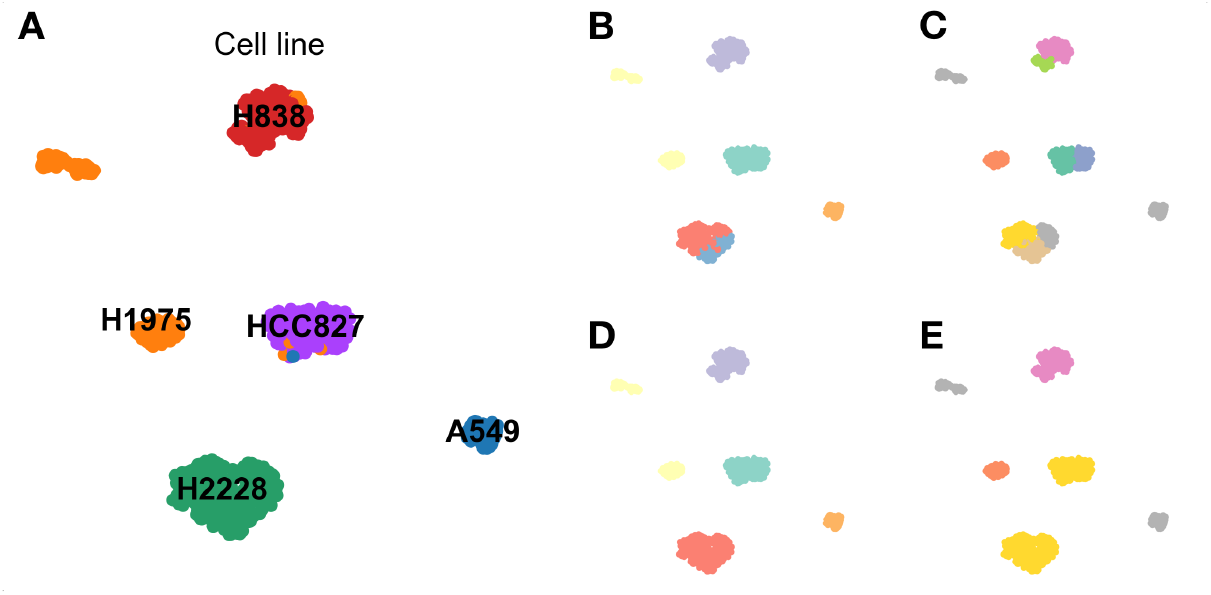
Analysis of a mixture of 5 known cell lines. (A) UMAP embedding coloured and labeled by cell line identity. H1975 cells (orange) show two distinct groups reflecting an internal heterogeneity. (B) UMAP embedding showing cells coloured by level 1 of the hierarchy proposed by the nested Stochastic Block Model. (C) UMAP embedding showing cells coloured according to the Leiden method at resolution *γ* = 1. (D) UMAP embedding coloured by the classification made by SCCAF when partitions in (B) are used. (E) UMAP embedding coloured by the classification made by SCCAF when partitions in (C) are used.

In all, these data support the suitability of `schist`, hence of Nested Stochastic Block Models, for cell group identification in single cell studies.

### Hierarchy modelling complies with biological properties

When grouping is performed by optimisation of modularity, there is often the implicit assumption that the resolution parameter reflects a hierarchical structure of the graph, that is communities are consistently grouped at lower resolutions. Not only this assumption is wrong, but it may also lead to spurious groupings in real experiments, whereas nSBM inherently encodes hierarchies by merging communities in a tree. The improper use of resolution parameter may lead to two types of errors: grouping of cells that are in fact distinct and creating an inconsistent hierarchy.

To show this we took advantage of public spatial RNA dataset of a coronal section of murine brain tissue profiled with 10X Visium H&E technology [48], as provided by the recently introduced package `SquidPy` [49]. We chose to stick to the given tissue annotation by the package authors. At default resolution, Leiden clustering resolves the tissue structure, as does the first level of the nSBM hierarchy (Fig. 3). When resolution is decreased (*e.g. γ* = 0.5), the dentate gyrus is incorrectly merged to the hippocampus, whereas nSBM correctly identifies the pyramidal layer.

**Figure 3:**
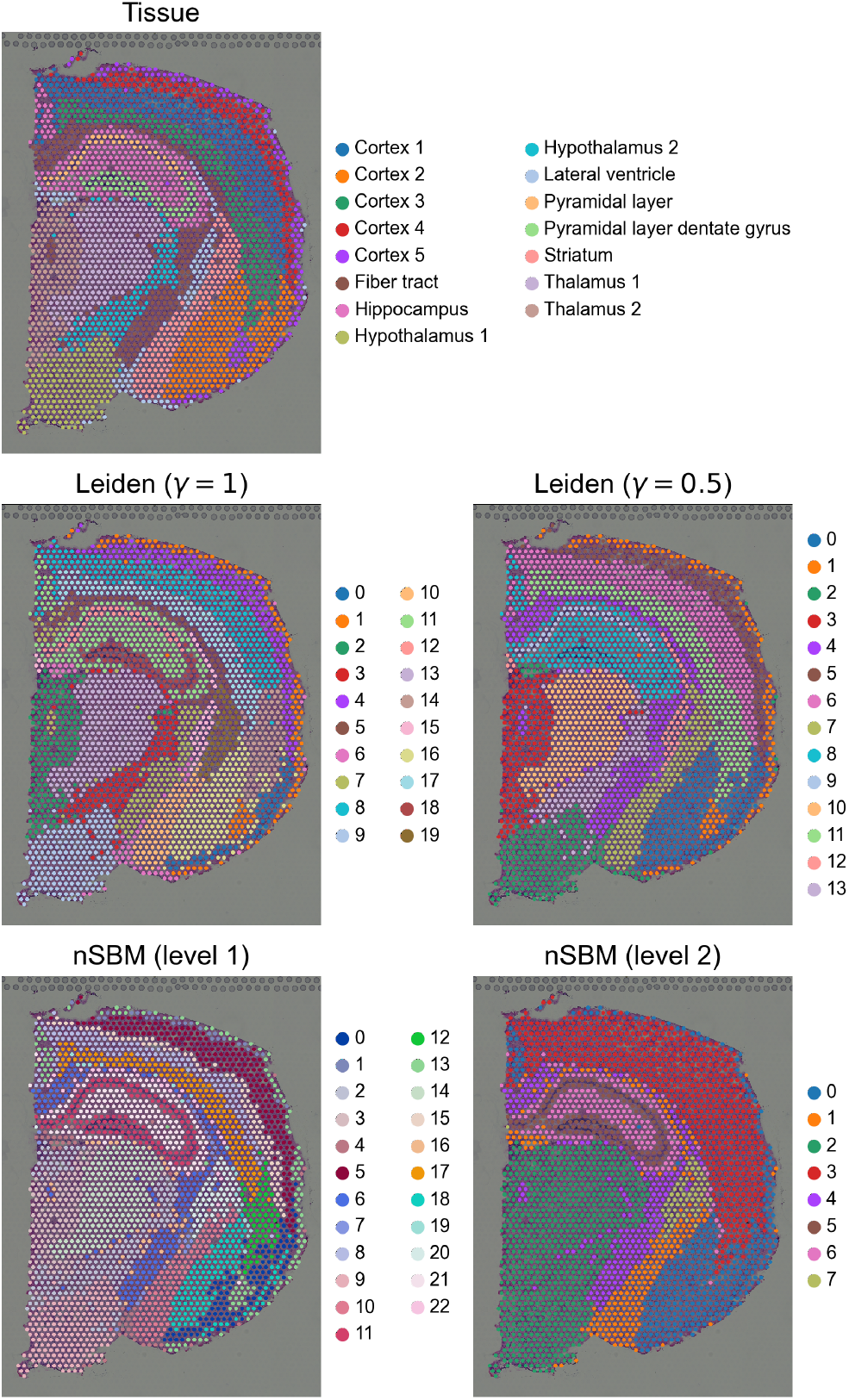
Analysis of spatial transcriptomics of a coronal section of mouse brain by Visium H&E. In the first panel, original tissue annotation is given. Tissues are well defined at default resolution for the standard approach. When resolution is decreased to *γ* = 0.5, cells from the dentate gyrus are merged to the hippocampus. When nSBM is used, the structure of the pyramidal is maintained at different levels of the hierarchy

In another context, we tested the effect on the interpretation of the hierarchy varying the resolution parameter. We analysed data for hematopoietic differentiation [50], previously used to benchmark the consistency of cell grouping with differentiation trajectories by graph abstraction [51] (Fig. S2A). Data show three major branchings (Erythroids, Neutrophils and Monocytes) stemming from the progenitor cells, mostly recapitulated by level 2 of the hierarchy computed by `schist` (Fig. 4). Not only the hierarchic model recapitulates the branching trajectories, also the cell groups appear to be consistent with the estimated pseudotime. Conversely, the Leiden method at default resolution identified 24 groups. By lowering the *γ* parameter we observed cell groups that merge and split at different resolutions disrupting the hierarchy (Fig. S3).

**Figure 4:**
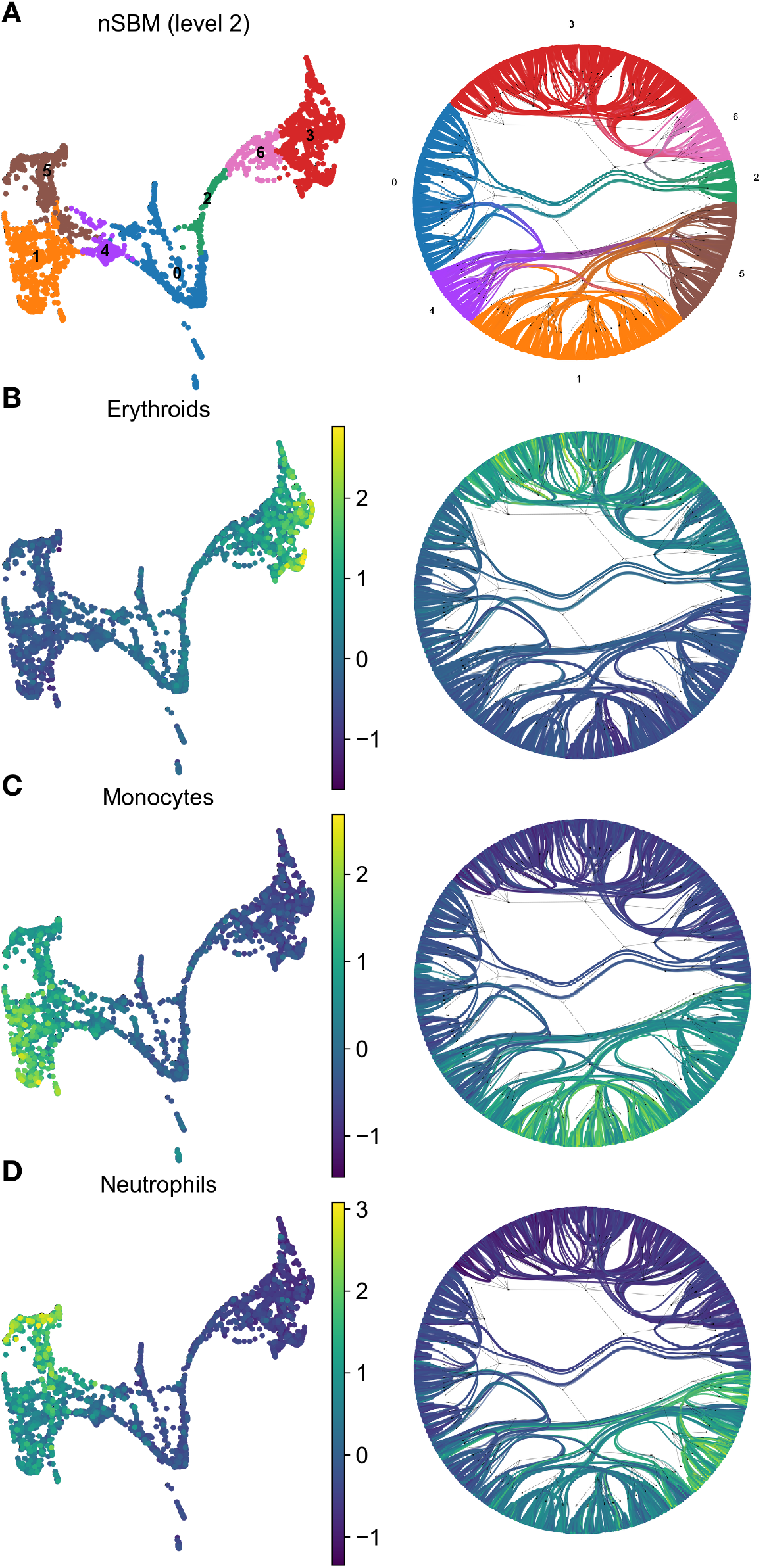
Analysis of hematopoietic differentiation. Each panel presents a low dimensional embedding of single cells next to a radial tree representation of the nSBM hierarchy. Cells are coloured according to groupings at level 2 of the hierarchy, group 0 marks the most primitive population (A). In subsequent panels, cells are coloured using a signature of Erythroid lineage (B), Monocytes (C) or Neutrophils (D).

In all, these data show that the common intuition that *γ* parameter acts as a thresholding factor over a hierarchy is wrong. Not only the hierarchy is not conserved, but also very different cell types may be mixed in spurious clusters. By using nSBM, `schist` is able to represent hierarchical relations in appropriate way. Moreover, the hierarchy appears to be more robust in aggregating different cell types at coarser scales.

### Cell marginals can be used to assess the data quality

By computing the consensus among multiple models, `schist` returns the marginal probability for each cell to belong to a specific cluster at each level of the hierarchy. Ideally, all cells should always be assigned with *p* = 1 to a cluster. When the uncertainty is maximal, cells are assigned to clusters randomly with *p* = 1/*B_i_*, where *B_i_* is the number of groups for the *i*-th level in the hierarchy. We sought to check if these probabilities could be interpreted in terms of data quality.

We devised a simple metric, *cell stability*, that is defined by the fraction of levels for which the marginal probability is higher than 1 — 1/*B_i_*. To do so, we only consider levels with at least two groups, hence excluding the root of the tree. We tested this metric on four datasets from [52] with different quality levels (iCELL8, MARS-seq, 10XV3 and Quartsz-seq2) (Fig. S4). By taking a summary metric, *e.g.* the mean 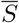 or the fraction of cells with *S* > 0.5, we observed that it correlates with the data quality (Table 1).

**Table 1:**
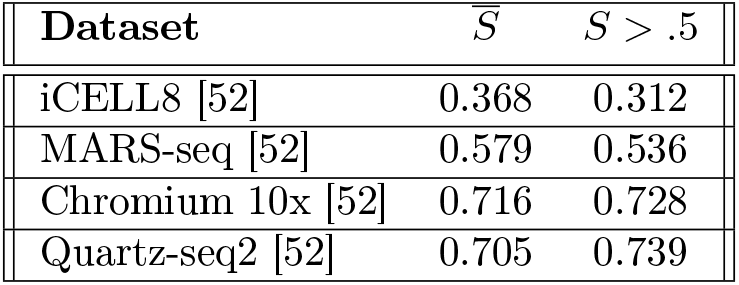
Cell stability as indicator of data quality. Table shows summary metrics derived from the Cell Stability calculated for various datasets. 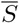 is the average Cell Stability over all cells, *S* > .5 indicates the fraction of cells with Cell Stability higher than 0.5.

These data suggest that measures of uncertainty of cell clustering can be useful for general quality control assessment. In addition to this, we foresee they could be used to isolate cells with specific patterns.

### Cell affinities can be used for label transfer

The modelling approach we adopted allows the estimation of the information required to describe a graph given any partitioning scheme, not limited to the solution given by the model itself. Differences in entropy can be used to perform model selection, that is we can choose which model better describes the data. We sought to exploit this property to address the task of annotating cells according to a reference sample. To this end we analysed datasets from [52], which includes mixtures of human PBMC and HEK293T cells profiled with various technologies. We chose cells profiled with 10X V3 platform as reference dataset and performed annotation on cells profiled with Quartz-seq2 or MARS-seq. These are at the extremes of the capability to distinguish cell types, so they provide good benchmark configurations for this task.

After preprocessing raw data according to the parameters given in [52], we integrated each dataset with 10XV3 into a unified representation using Harmony [54], and computed the *k*NN graph. In each merged dataset, we retained cell type annotations for 10X cells, while we assigned a “Unknown” label to all cells derived from the other technology (i.e. MARS-seq or Quartz-seq2). We then calculated the *cell affinity* matrix, that is we computed the difference in entropy that can be observed by assigning each cell to each annotation cluster, this being either one of the original cell types or “Unknown”. Once the matrix has been computed, each cell from the query data is assigned to the group with the highest likelihood. The rationale behind this approach is that if cells belong to the same annotation group, then more information is required to describe the graph if they were annotated as different cell types; hence, cells from the query datasets should retain their “Unknown” label if and only if there is not enough evidence to associate them to another group. We compared the accuracy of the outcome to *k*NN classification, given by the closest entry in the *k*NN graph, and to `ingest`, a tool included in `scanpy` based on *k*NN classification of UMAP embeddings. Analysis of a well defined dataset, such as Quartz-seq2, reveals that the three approaches are equally good in classifying unknown cells (Fig. 5, central column), with accuracies ranging from .870 to .927. When data are noisy, instead, *k*NN-based methods show low accuracy and a tendency to assign the most represented cell group (HEK293T) to the unlabelled cells. This misannotation is particular evident for `ingest`, in which only CD4 T cells and HEK cells are transferred, resulting in the lowest accuracy (0.243). Conversely, `schist` is able to assign correct labels with higher accuracy (0.641). Moreover, *k*NN methods assign a label to each cell, whereas `schist` does not relabel cells if there are no sufficient evidence (e.g. the “Unknown” state is the most likely). Interestingly, we found that for the largest part of cells without assigned label, the second choice by affinity ranking was indeed the appropriate one (Fig. S5).

**Figure 5:**
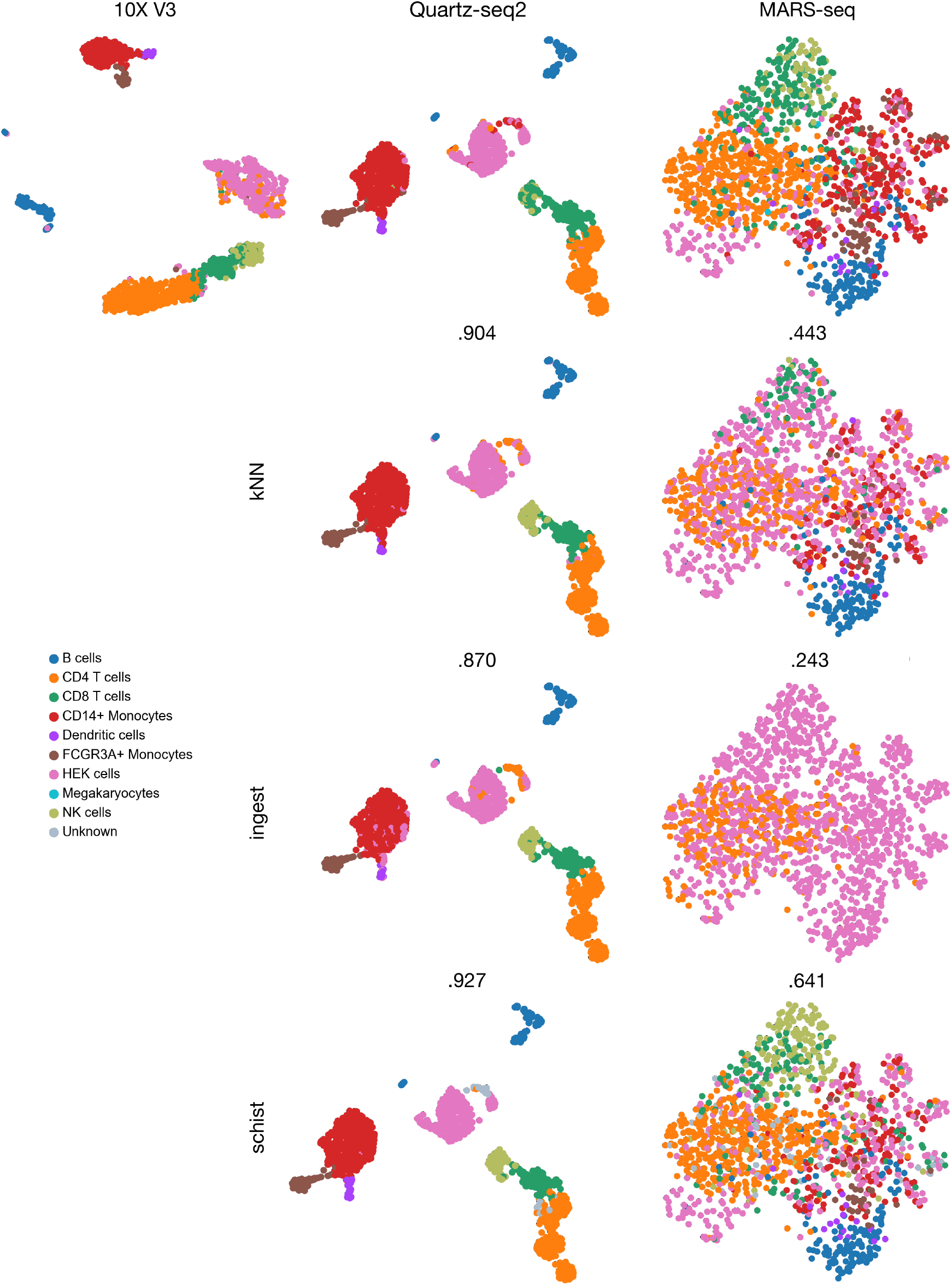
Label transfer using SBM. The first line reports UMAP embeddings for datasets profiled with Chromium 10X V3, Quartz-seq2 and MARS-seq, each annotated by known cell types. Quartz-seq2 and MARS-seq were reannotated using *k*NN method, `sc.tl.ingest()` or `schist`. The accuracy of each label transfer task is reported above the corresponding UMAP.

### Choice of an optimal hierarchy level

The Nested Stochastic Block Model, hence `schist`, fits a hierarchical model of communities into a graph. When it comes to analysis of single cell data, it means that the cells are best described by the hierarchy itself and that cells *can* be grouped consistently at each level of the tree. In addition, the size of groups at the deepest level scales as *O*(*N*/log *N*) [44], where *N* is the number of cells. Given the current throughput in single cell experiments (~10k cells), the number of groups becomes intractable. For this reason, in most of single cell experiments, it is preferable to identify an optimal level of the hierarchy that best resembles the cell properties at the scale they can be validated.

A possible strategy is based on Random Matrix Theory, as suggested by the authors of the SC3 package [15], for which a suitable number of clusters, 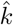, is determined by the number of eigenvalues of the 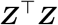 matrix (where ***Z*** is the normalised count matrix) significantly different at *p* < .001 from the appropriate Tracy-Widom distribution. According to this strategy, the optimal level *i^k^* is the one that minimises the number of partitions and 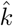:

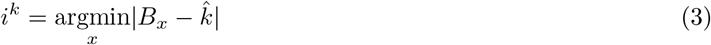

where *B_x_* is the number of non empty partitions at level *x*.

An alternative strategy is to evaluate the behaviour of modularity at different hierarchy levels. While nSBM does not optimise the graph modularity *Q*, we observed that this tends to be maximal for the level better describing known cell populations, so the optimal level *i^Q^* is

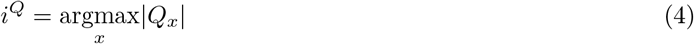

Where *Q_x_* is modularity at level *x*. We collected values arising from both the approaches for some datasets used in this work (Table 2 and Fig. S6)

**Table 2:**
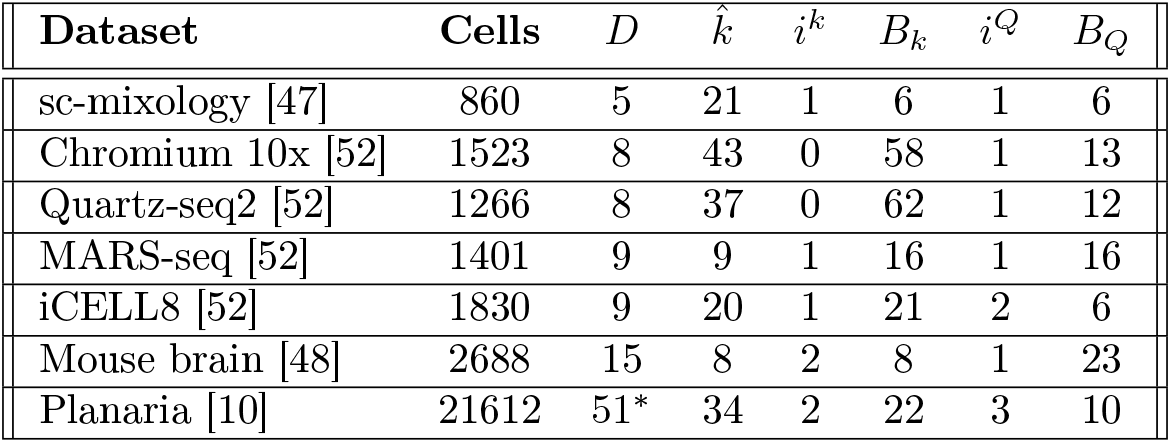
Selection of the optimal level in the nSBM hierarchy. For each dataset we report the number of groups *D* that were given by the authors. The optimal level selection should recover a number of groups in the order of magnitude of *D*. Value of *D* in Planaria dataset is derived from manual curation of Louvain clustering. 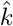: number of groups according to RMT, *i^k^*: level selected according to RMT criterion, *B_k_*: number of partitions at level *i^k^*, *i^Q^*: level at which modularity is maximal, *B_Q_*: number of groups at level *i^Q^*

| Dataset | Cells | $D$ | $\hat{k}$ | $i^k$ | $B_k$ | $i^Q$ | $B_Q$ |
| --- | --- | --- | --- | --- | --- | --- | --- |
| sc-mixology [47] | 860 | 5 | 21 | 1 | 6 | 1 | 6 |
| Chromium 10x [52] | 1523 | 8 | 43 | 0 | 58 | 1 | 13 |
| Quartz-seq2 [52] | 1266 | 8 | 37 | 0 | 62 | 1 | 12 |
| MARS-seq [52] | 1401 | 9 | 9 | 1 | 16 | 1 | 16 |
| iCELL8 [52] | 1830 | 9 | 20 | 1 | 21 | 2 | 6 |
| Mouse brain [48] | 2688 | 15 | 8 | 2 | 8 | 1 | 23 |
| Planaria [10] | 21612 | 51* | 34 | 2 | 22 | 3 | 10 |

As expected, the larger the network, the higher the optimal level. For relatively small datasets (*i.e.* less than 10k cells), the first level of the hierarchy contains a number of groups in line with how many observable populations are.

### Analysis of runtimes

Minimisation of the nSBM is a process that requires a large amount of computational resources. While the underlying `graph-tool` library is efficient in exploring the solution space using a multiflip MCMC sampling strategy, the number of required iterations before convergence is considerable and the running time scales linearly with the number of edges. Moreover, to collect a consensus partition, we minimise multiple models (default: 100) that need to be averaged. To give a reference, we report runtimes for some example datasets in Table 3 on a commodity hardware (Intel i7@2.8 GHz, 32 GB RAM). Compared to Leiden approach, nSBM requires at best ~ 6× times more, and ~ 30× at worst. A reasonably fast alternative to the nSBM is the Planted Partition Block Model (PPBM), for which we also report runtimes. The PPBM [46] is able to find statistically significant assortative modules and eliminates the resolution parameter; differently from nSBM, PPBM is not hierarchic.

**Table 3:**
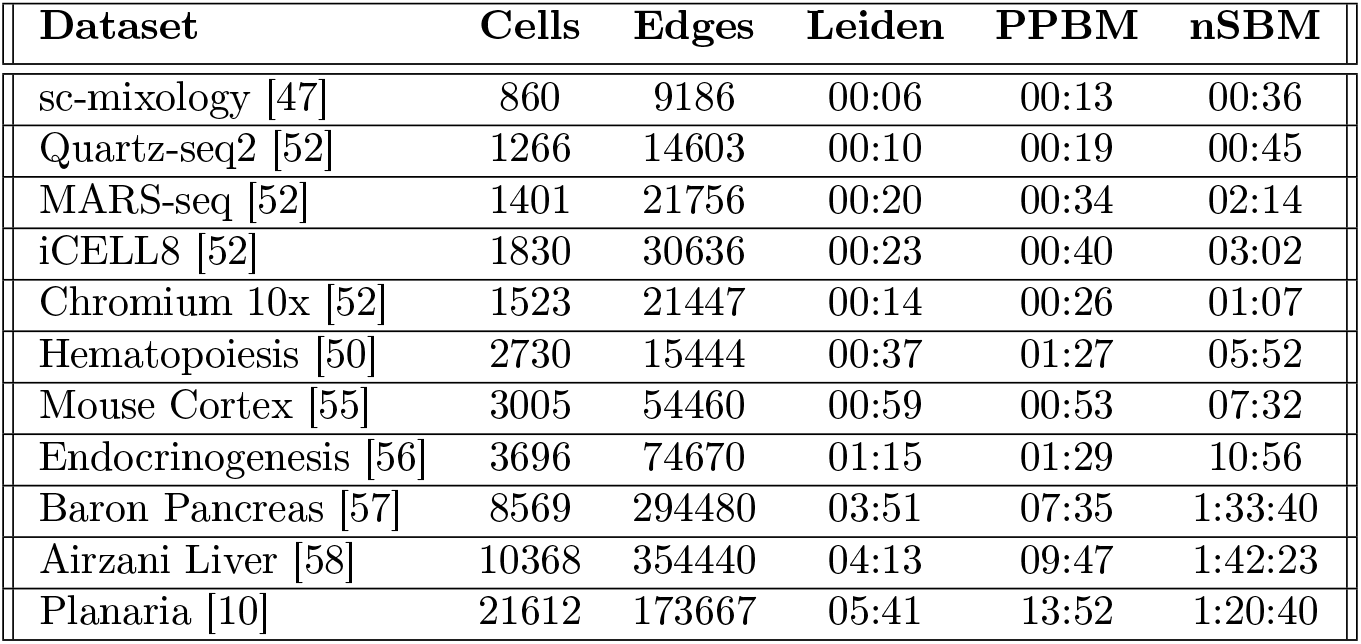
Time required to run different partitioning strategies implemented in `schist` on various datasets. All approaches fit 100 models. Number of nodes and edges refer to the structure of the *k*NN graph as built by `scanpy`. Times are expressed in MM:SS.

| Dataset | Cells | Edges | Leiden | PPBM | nSBM |
| --- | --- | --- | --- | --- | --- |
| sc-mixology [47] | 860 | 9186 | 00:06 | 00:13 | 00:36 |
| Quartz-seq2 [52] | 1266 | 14603 | 00:10 | 00:19 | 00:45 |
| MARS-seq [52] | 1401 | 21756 | 00:20 | 00:34 | 02:14 |
| iCELL8 [52] | 1830 | 30636 | 00:23 | 00:40 | 03:02 |
| Chromium 10x [52] | 1523 | 21447 | 00:14 | 00:26 | 01:07 |
| Hematopoiesis [50] | 2730 | 15444 | 00:37 | 01:27 | 05:52 |
| Mouse Cortex [55] | 3005 | 54460 | 00:59 | 00:53 | 07:32 |
| Endocrinogenesis [56] | 3696 | 74670 | 01:15 | 01:29 | 10:56 |
| Baron Pancreas [57] | 8569 | 294480 | 03:51 | 07:35 | 1:33:40 |
| Airzani Liver [58] | 10368 | 354440 | 04:13 | 09:47 | 1:42:23 |
| Planaria [10] | 21612 | 173667 | 05:41 | 13:52 | 1:20:40 |

## Conclusions

Identification of cells sharing similar properties in single cell experiments is of paramount importance. A large number of approaches have been described, although the standardisation of analysis pipelines converged to methods that are based on modularity optimisation. We tackled the biological problem using a different approach, nSBM, which has several advantages over existing techniques. The most important advantage is the hierarchical definition of cell groups which eliminates the choice of an arbitrary threshold on clustering resolution. In addition, we showed that the hierarchy itself could have a biological interpretation, implying that the hierarchical model is a valid representation of the cell ensemble.

The Bayesian formulation of Stochastic Block Models provides the possibility to perform inference on a graph for any partition configuration, thus allowing reliable model selection using an interpretable measure, entropy. We exploited this property to perform label transfer with high accuracy and with the possibility to discard cells with unreliable assignments. In all, `schist` facilitates the adoption of nSBM by the bioinformatics community and exposes a robust framework to perform tasks that go beyond the principled identification of cell clusters.

The major drawback of adopting this strategy is the substantial increase of runtimes. As observed, model minimisation may be hundred times slower than the extremely fast Leiden approach. It should be noted that `schist` initialises multiple models that are treated by multiple concurrent processes. `graph-tool` itself supports CPU-level parallelisation for some of its tasks. These optimisations are well suited for clustered computing infrastructure. Further development, possibly including GPU-level parallelisation, is surely required to accomodate the large size of datasets that are being produced.

## Materials and Methods

Unless differently stated, all the analysis were produced using `scanpy` v1.7.1 [22] and `schist` v0.7.6 and the corresponding dependencies. All models were initialised 100 times, herein including Leiden partitioning for which we also calculated the consensus partition.

### Analysis of Random data

Data were retrieved in `scanpy` environment using `sc.datasets.pbmc3k_processed()` function. The random *k*NN graph was obtained shuffling the node labels of each edge. UMAP embedding was recomputed after randomisation using the shuffled graph.

### Analysis of cell mixtures

Data and metadata for five cell mixture profiled by Chromium 10x were downloaded from the sc-mixology repository (https://github.com/LuyiTian/sc_mixology). Cells with less than 200 genes were excluded, as genes detected in less than 3 cells. Cells with less than 5% of mitochondrial genes were retained for subsequent analysis. Data were normalised and log-transformed; number of genes and percentage of mitochondrial genes were regressed out. *k*NN graph was built with default parameters (50 components and 15 nearest neighbours).Data were assessed by SCCAF using cell line annotation. Mean cross-validated accuracy was set as target for all the models.

### Analysis of Visium H&E data

Data were retrieved using `squidpy.datasets.visium_hne_adata()` built-in function, without further processing. Leiden clustering was performed using `schist.inference.leiden()` function, allowing for 100 initialisations, with resolutions *γ* = 1 and *γ* = 0.5.

### Analysis of hematopoietic differentiation

Data were retrieved using `scanpy`’s built-in functions and were processed as in [51], except for *k*NN graph built using 30 principal components, 30 neighbours and diffmap as embedding. Gene signatures were calculated using the following gene lists

1. Erythroids: Gata1, Klf1, Epor, Gypa, Hba-a2, Hba-a1, Spi1
2. Neutrophils, Elane, Cebpe, Ctsg, Mpo, Gfi1
3. Monocytes, Irf8, Csf1r, Ctsg, Mpo

### Processing of PBMC data from various platforms

Count matrices were downloaded from GEO using the following accession numbers: GSE133535 (Chromium 10Xv3), GSE133543 (Quartz-seq2), GSE133542 (MARS-seq) and GSE133541 (iCELL8). Data were processed according to the methods in the original paper [52]. Briefly, cells with less than 10,000 total number of reads as well as the cells having less than 65% of the reads mapped to their reference genome were discarded. Cells in the 95th percentile of the number of genes/cell and those having less than 25% mitochondrial gene content were included in the downstream analyses. Genes that were expressed in less than five cells were removed. Data were normalised and log-transformed, highly variable genes were detected at minimal dispersion equal to 0.5. Neighbourhood graph was built using 30 principal components and 20 neighbours.

### Label transfer

Processed data for MARS-seq or Quart-seq2 platforms were merged to data for 10X V3. Merged data were then processed using Harmony [54] by the `scanpy.external.pp.harmony_integrate()` function with default parameters. Cells not belonging to the 10X data were assigned an “Unknown” label. We calculated cell affinity to each annotation label using `schist.tl.calculated_affinity()` function. We assigned the most affine annotation only to “Unknown” cells. For *k*NN-based procedure, we built a *k*NN graph on the merged data using `pynndescent` library on the 10XV3 subset of cells in the merged data, then we assigned “Unknown” cells to the closest entry in the graph. Assignment by `scanpy.tl.ingest()` was performed using default parameters.

## Acknowledgements

We would like to thank Tiago de Paula Peixoto (Central European University, ISI Foundation) and Giovanni Petri (ISI Foundation) for the discussions and the precious hints. We also would like to thank all people at COSR, in particular Giovanni Tonon and Paolo Provero. This work has been supported by Accelerator Award: A26815 entitled: “Single-cell cancer evolution in the clinic” funded through a partnership between Cancer Research UK and Fondazione AIRC.

## References

[1] Svensson V, Vento-Tormo R, Teichmann SA. Exponential scaling of single-cell RNA-seq in the past decade. Nature Protocols. 2018;13(4):599–604. doi:10.1038/nprot.2017.149.

[2] Guo J, Grow EJ, Mlcochova H, Maher GJ, Lindskog C, Nie X, et al. The adult human testis transcriptional cell atlas. Cell Research. 2018;28(12):1141–1157. doi:10.1038/s41422-018-0099-2.

[3] Vento-Tormo R, Efremova M, Botting RA, Turco MY, Vento-Tormo M, Meyer KB, et al. Singlecell reconstruction of the early maternal-fetal interface in humans. Nature. 2018;563(7731):347–353. doi:10.1038/s41586-018-0698-6.

[4] Rozenblatt-Rosen O, Regev A, Oberdoerffer P, Nawy T, Hupalowska A, Rood JE, et al. The Human Tumor Atlas Network: Charting Tumor Transitions across Space and Time at Single-Cell Resolution. Cell. 2020;181(2):236–249. doi:10.1016/j.cell.2020.03.053.

[5] Tirosh I, Izar B, Prakadan SM, Wadsworth MH, Treacy D, Trombetta JJ, et al. Dissecting the multicellular ecosystem of metastatic melanoma by single-cell RNA-seq. Science. 2016;352(6282):189–196. doi:10.1126/science.aad0501.

[6] Patel AP, Tirosh I, Trombetta JJ, Shalek AK, Gillespie SM, Wakimoto H, et al. Single-cell RNA-seq highlights intratumoral heterogeneity in primary glioblastoma. Science. 2014;344(6190):1396–1401. doi:10.1126/science.1254257.

[7] Neftel C, Laffy J, Filbin MG, Hara T, Shore ME, Rahme GJ, et al. An integrative model of cellular states, plasticity, and genetics for glioblastoma. Cell. 2019;178(4):835–849.e21. doi:10.1016/j.cell.2019.06.024.

[8] Rosenberg AB, Roco CM, Muscat RA, Kuchina A, Sample P, Yao Z, et al. Single-cell profiling of the developing mouse brain and spinal cord with split-pool barcoding. Science. 2018;360(6385):176–182. doi:10.1126/science.aam8999.

[9] Wagner DE, Weinreb C, Collins ZM, Briggs JA, Megason SG, Klein AM. Single-cell mapping of gene expression landscapes and lineage in the zebrafish embryo. Science. 2018;360(6392):981–987. doi:10.1126/science.aar4362.

[10] Plass M, Solana J, Wolf FA, Ayoub S, Misios A, Glažar P, et al. Cell type atlas and lineage tree of a whole complex animal by single-cell transcriptomics. Science. 2018;360(6391). doi:10.1126/science.aaq1723.

[11] Regev A, Teichmann SA, Lander ES, Amit I, Benoist C, Birney E, et al. The human cell atlas. eLife. 2017;6. doi:10.7554/eLife.27041.

[12] Wang B, Zhu J, Pierson E, Ramazzotti D, Batzoglou S. Visualization and analysis of single-cell RNA-seq data by kernel-based similarity learning. Nature Methods. 2017;14(4):414–416. doi:10.1038/nmeth.4207.

[13] Lin P, Troup M, Ho JWK. CIDR: Ultrafast and accurate clustering through imputation for single-cell RNA-seq data. Genome Biology. 2017;18(1):59. doi:10.1186/s13059-017-1188-0.

[14] Huh R, Yang Y, Jiang Y, Shen Y, Li Y. SAME-clustering: Single-cell Aggregated Clustering via Mixture Model Ensemble. Nucleic Acids Research. 2020;48(1):86–95. doi:10.1093/nar/gkz959.

[15] Kiselev VY, Kirschner K, Schaub MT, Andrews T, Yiu A, Chandra T, et al. SC3: consensus clustering of single-cell RNA-seq data. Nature Methods. 2017;14(5):483–486. doi:10.1038/nmeth.4236.

[16] Ranjan B, Schmidt F, Sun W, Park J, Honardoost MA, Tan J, et al. scConsensus: combining supervised and unsupervised clustering for cell type identification in single-cell RNA sequencing data. BMC Bioinformatics. 2021;22(1):186. doi:10.1186/s12859-021-04028-4.

[17] Li X, Wang K, Lyu Y, Pan H, Zhang J, Stambolian D, et al. Deep learning enables accurate clustering with batch effect removal in single-cell RNA-seq analysis. Nature Communications. 2020;11(1):2338. doi:10.1038/s41467-020-15851-3.

[18] Krzak M, Raykov Y, Boukouvalas A, Cutillo L, Angelini C. Benchmark and Parameter Sensitivity Analysis of Single-Cell RNA Sequencing Clustering Methods. Frontiers in genetics. 2019;10:1253. doi:10.3389/fgene.2019.01253.

[19] Kiselev VY, Andrews TS, Hemberg M. Challenges in unsupervised clustering of single-cell RNA-seq data. Nature Reviews Genetics. 2019;20(5):273–282. doi:10.1038/s41576-018-0088-9.

[20] Duò A, Robinson MD, Soneson C. A systematic performance evaluation of clustering methods for single-cell RNA-seq data. F1000Research. 2018;7:1141. doi:10.12688/f1000research.15666.2.

[21] Butler A, Hoffman P, Smibert P, Papalexi E, Satija R. Integrating single-cell transcriptomic data across different conditions, technologies, and species. Nature Biotechnology. 2018;36(5):411–420. doi:10.1038/nbt.4096.

[22] Wolf FA, Angerer P, Theis FJ. SCANPY: large-scale single-cell gene expression data analysis. Genome Biology. 2018;19(1):15. doi:10.1186/s13059-017-1382-0.

[23] Setty M, Kiseliovas V, Levine J, Gayoso A, Mazutis L, Pe’er D. Characterization of cell fate probabilities in single-cell data with Palantir. Nature Biotechnology. 2019;37(4):451–460. doi:10.1038/s41587-019-0068-4.

[24] Lange M, Bergen V, Klein M, Setty M, Reuter B, Bakhti M, et al. CellRank for directed single-cell fate mapping. BioRxiv. 2020;doi:10.1101/2020.10.19.345983.

[25] Bergen V, Lange M, Peidli S, Wolf FA, Theis FJ. Generalizing RNA velocity to transient cell states through dynamical modeling. Nature Biotechnology. 2020;38(12):1408–1414. doi:10.1038/s41587-020-0591-3.

[26] Blondel VD, Guillaume JL, Lambiotte R, Lefebvre E. Fast unfolding of communities in large networks. Journal of Statistical Mechanics: Theory and Experiment. 2008;2008(10):P10008. doi:10.1088/1742-5468/2008/10/P10008.

[27] Traag VA, Waltman L, van Eck NJ. From Louvain to Leiden: guaranteeing well-connected communities. Scientific Reports. 2019;9(1):5233. doi:10.1038/s41598-019-41695-z.

[28] Levine JH, Simonds EF, Bendall SC, Davis KL, Amir EaD, Tadmor MD, et al. Data-Driven Phenotypic Dissection of AML Reveals Progenitor-like Cells that Correlate with Prognosis. Cell. 2015;162(1):184–197. doi:10.1016/j.cell.2015.05.047.

[29] Newman MEJ, Girvan M. Finding and evaluating community structure in networks. Physical Review E, Statistical, Nonlinear, and Soft Matter Physics. 2004;69(2 Pt 2):026113. doi:10.1103/PhysRevE.69.026113.

[30] Traag VA, Van Dooren P, Nesterov Y. Narrow scope for resolution-limit-free community detection. Physical Review E. 2011;84(1). doi:10.1103/PhysRevE.84.016114.

[31] Reichardt J, Bornholdt S. Statistical mechanics of community detection. Physical Review E. 2006;74(1). doi:10.1103/PhysRevE.74.016110.

[32] Lähnemann D, Köster J, Szczurek E, McCarthy DJ, Hicks SC, Robinson MD, et al. Eleven grand challenges in single-cell data science. Genome Biology. 2020;21(1):31. doi:10.1186/s13059-020-1926-6.

[33] Fortunato S, Barthélemy M. Resolution limit in community detection. Proceedings of the National Academy of Sciences of the United States of America. 2007;104(1):36–41. doi:10.1073/pnas.0605965104.

[34] Guimerà R, Sales-Pardo M, Amaral LAN. Modularity from fluctuations in random graphs and complex networks. Physical Review E. 2004;70(2). doi:10.1103/PhysRevE.70.025101.

[35] Baran Y, Bercovich A, Sebe-Pedros A, Lubling Y, Giladi A, Chomsky E, et al. MetaCell: analysis of singlecell RNA-seq data using K-nn graph partitions. Genome Biology. 2019;20(1):206. doi:10.1186/s13059-019-1812-2.

[36] Tang M, Kaymaz Y, Logeman BL, Eichhorn S, Liang ZS, Dulac C, et al. Evaluating Single-Cell Cluster Stability Using The Jaccard Similarity Index. Bioinformatics. 2020;doi:10.1093/bioinformatics/btaa956.

[37] Xu C, Su Z. Identification of cell types from single-cell transcriptomes using a novel clustering method. Bioinformatics. 2015;31(12):1974–1980. doi:10.1093/bioinformatics/btv088.

[38] Miao Z, Moreno P, Huang N, Papatheodorou I, Brazma A, Teichmann SA. Putative cell type discovery from single-cell gene expression data. Nature Methods. 2020;17(6):621–628. doi:10.1038/s41592-020-0825-9.

[39] Holland PW, Laskey KB, Leinhardt S. Stochastic blockmodels: First steps. Social networks. 1983;5(2):109–137. doi:10.1016/0378-8733(83)90021-7.

[40] Peixoto TP. Nonparametric Bayesian inference of the microcanonical stochastic block model. Physical review E. 2017;95(1-1):012317. doi:10.1103/PhysRevE.95.012317.

[41] Karrer B, Newman MEJ. Stochastic blockmodels and community structure in networks. Physical Review E, Statistical, Nonlinear, and Soft Matter Physics. 2011;83(1 Pt 2):016107. doi:10.1103/PhysRevE.83.016107.

[42] Peixoto TP. Parsimonious module inference in large networks. Physical Review Letters. 2013;110(14):148701. doi:10.1103/PhysRevLett.110.148701.

[43] Peixoto TP. Efficient Monte Carlo and greedy heuristic for the inference of stochastic block models. Physical Review E, Statistical, Nonlinear, and Soft Matter Physics. 2014;89(1):012804. doi:10.1103/PhysRevE.89.012804.

[44] Peixoto TP. Hierarchical Block Structures and High-Resolution Model Selection in Large Networks. Physical Review X. 2014;4(1):011047. doi:10.1103/PhysRevX.4.011047.

[45] Peixoto TP. Revealing Consensus and Dissensus between Network Partitions. Physical Review X. 2021;11(2):021003. doi:10.1103/PhysRevX.11.021003.

[46] Zhang L, Peixoto TP. Statistical inference of assortative community structures. Physical Review Research. 2020;2(4):043271. doi:10.1103/PhysRevResearch.2.043271.

[47] Tian L, Dong X, Freytag S, Lê Cao KA, Su S, JalalAbadi A, et al. Benchmarking single cell RNA-sequencing analysis pipelines using mixture control experiments. Nature Methods. 2019;16(6):479–487. doi:10.1038/s41592-019-0425-8.

[48] Gracia Villacampa E, Larsson L, Kvastad L, Andersson A, Carlson J, Lundeberg J. Genome-wide Spatial Expression Profiling in FFPE Tissues. BioRxiv. 2020;doi:10.1101/2020.07.24.219758.

[49] Palla G, Spitzer H, Klein M, Fischer DS, Schaar AC, Kuemmerle LB, et al. Squidpy: a scalable framework for spatial single cell analysis. BioRxiv. 2021;doi:10.1101/2021.02.19.431994.

[50] Paul F, Arkin Y, Giladi A, Jaitin DA, Kenigsberg E, Keren-Shaul H, et al. Transcriptional heterogeneity and lineage commitment in myeloid progenitors. Cell. 2015;163(7):1663–1677. doi:10.1016/j.cell.2015.11.013.

[51] Wolf FA, Hamey FK, Plass M, Solana J, Dahlin JS, Göttgens B, et al. PAGA: graph abstraction reconciles clustering with trajectory inference through a topology preserving map of single cells. Genome Biology. 2019;20(1):59. doi:10.1186/s13059-019-1663-x.

[52] Mereu E, Lafzi A, Moutinho C, Ziegenhain C, McCarthy DJ, Álvarez-Varela A, et al. Benchmarking single-cell RNA-sequencing protocols for cell atlas projects. Nature Biotechnology. 2020;38(6):747–755. doi:10.1038/s41587-020-0469-4.

[53] Zappia L, Phipson B, Oshlack A. Splatter: simulation of single-cell RNA sequencing data. Genome Biology. 2017;18(1):174. doi:10.1186/s13059-017-1305-0.

[54] Korsunsky I, Millard N, Fan J, Slowikowski K, Zhang F, Wei K, et al. Fast, sensitive and accurate integration of single-cell data with Harmony. Nature Methods. 2019;16(12):1289–1296. doi:10.1038/s41592-019-0619-0.

[55] Zeisel A, Muñoz-Manchado AB, Codeluppi S, Lönnerberg P, La Manno G, Juréus A, et al. Brain structure. Cell types in the mouse cortex and hippocampus revealed by single-cell RNA-seq. Science. 2015;347(6226):1138–1142. doi:10.1126/science.aaa1934.

[56] Bastidas-Ponce A, Tritschler S, Dony L, Scheibner K, Tarquis-Medina M, Salinno C, et al. Comprehensive single cell mRNA profiling reveals a detailed roadmap for pancreatic endocrinogenesis. Development. 2019;146(12). doi:10.1242/dev.173849.

[57] Baron M, Veres A, Wolock SL, Faust AL, Gaujoux R, Vetere A, et al. A Single-Cell Transcriptomic Map of the Human and Mouse Pancreas Reveals Inter- and Intra-cell Population Structure. Cell Systems. 2016;3(4):346–360.e4. doi:10.1016/j.cels.2016.08.011.

[58] Aizarani N, Saviano A, Sagar, Mailly L, Durand S, Herman JS, et al. A human liver cell atlas reveals heterogeneity and epithelial progenitors. Nature. 2019;572(7768):199–204. doi:10.1038/s41586-019-1373-2.

