## Supplementary figures for "Nested Stochastic Block Models Applied to the Analysis of Single Cell Data"

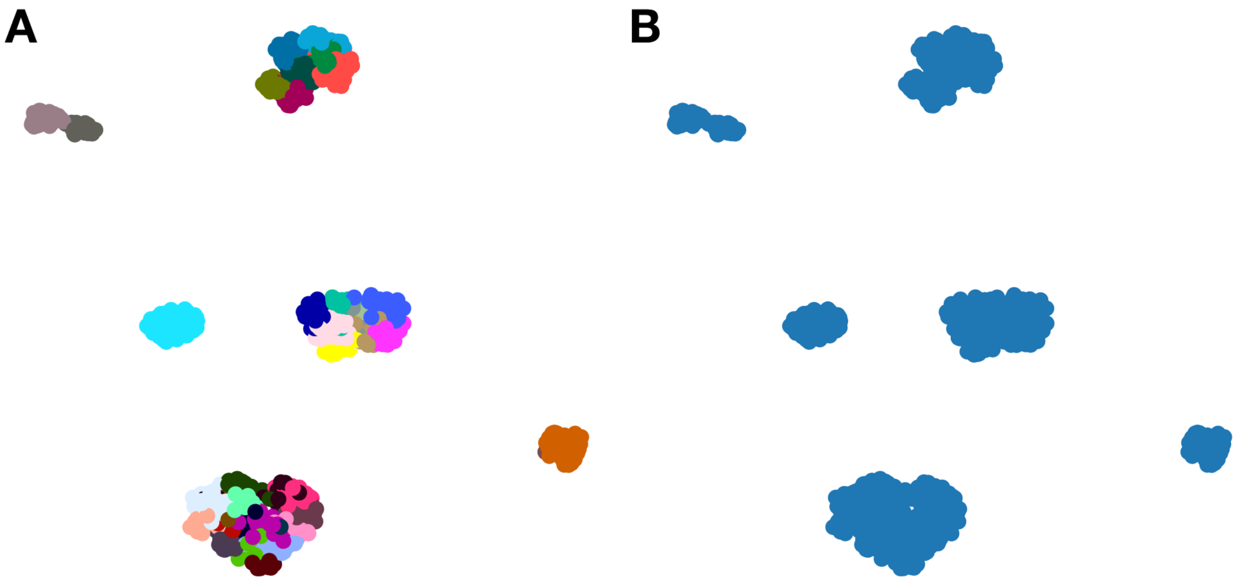


FIGURE S1: Analysis of a mixture of 5 known cell lines. (A) UMAP embedding of cell lines from the sc-mixology experiment coloured by the level 0 of the hierarchy proposed by the nested Stochastic Block Model (B) UMAP embedding coloured by the classification made by SCCAF when partitions in (A) are used.

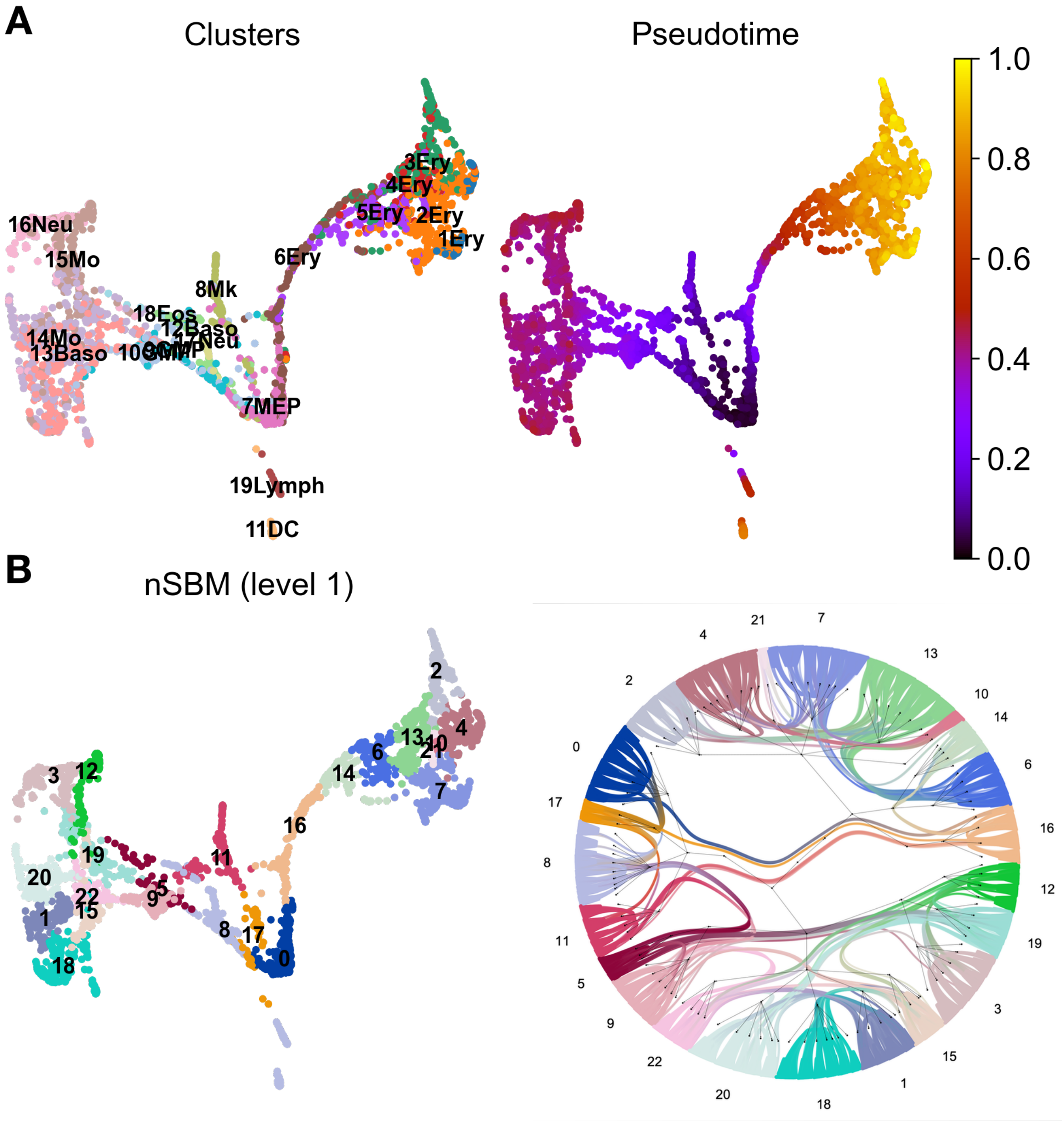


FIGURE S2: Analysis of hematopoietic differentiation. (A) Low dimensional embedding of single cells coloured by original cell type and pseudotime. (B) Cells are coloured according to the nSBM grouping at level 3 of the hierarchy, next to a radial tree representation of the same model.

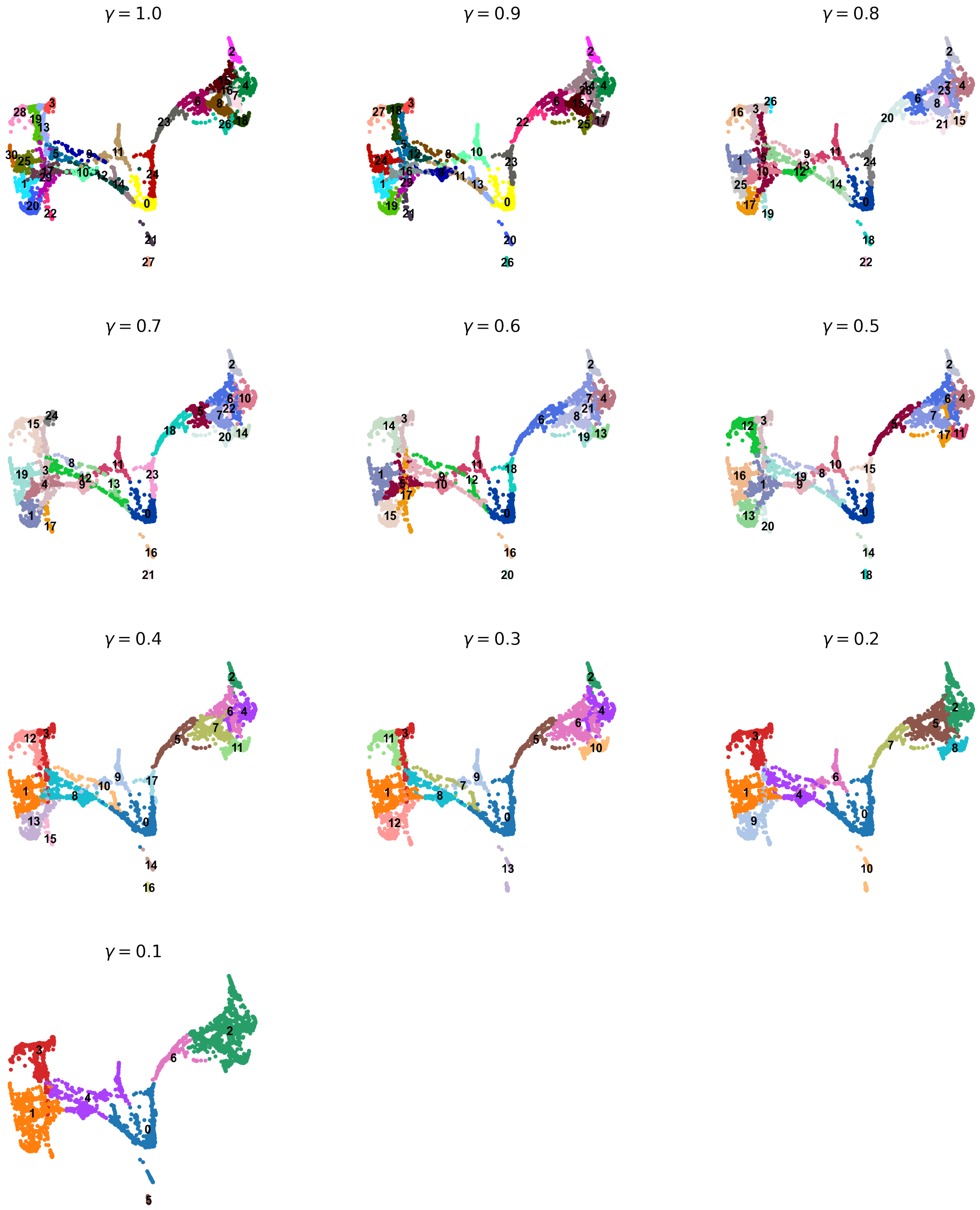


Figure S3: Low dimension embedding of single cells for hematopoietic differentiation coloured according to Leiden clustering at decreasing resolution, from 1.0 to 0.1. Lowering the distribution does not grant that cells are grouped in a hierarchical way, *e.g.* groups 6 and 8 in the Erythroid branch at resolution $\gamma = 1$ are merged or split at coarser resolutions ($\gamma = 0.6$ and $\gamma = 0.3$).

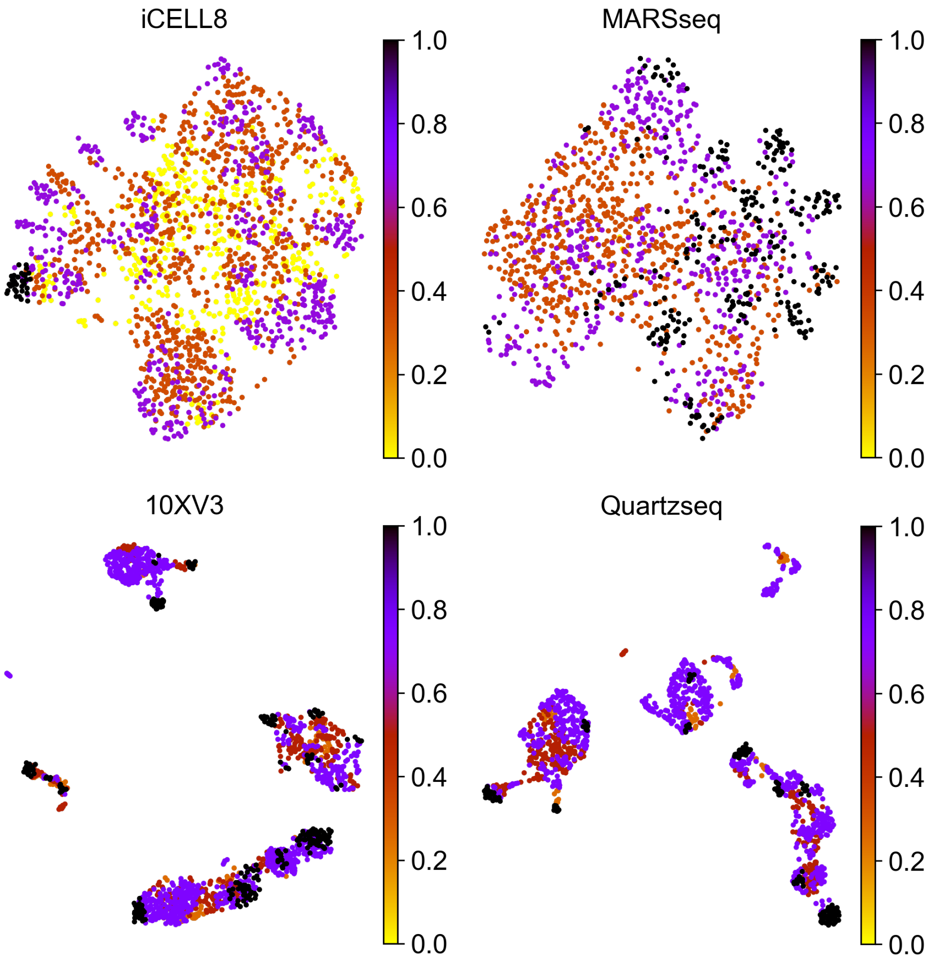


FIGURE S4: Cell Stability. UMAP embeddings of PBMC data profiled with four different technologies ranked by their quality. Cells are coloured by the Cell Stability metric.

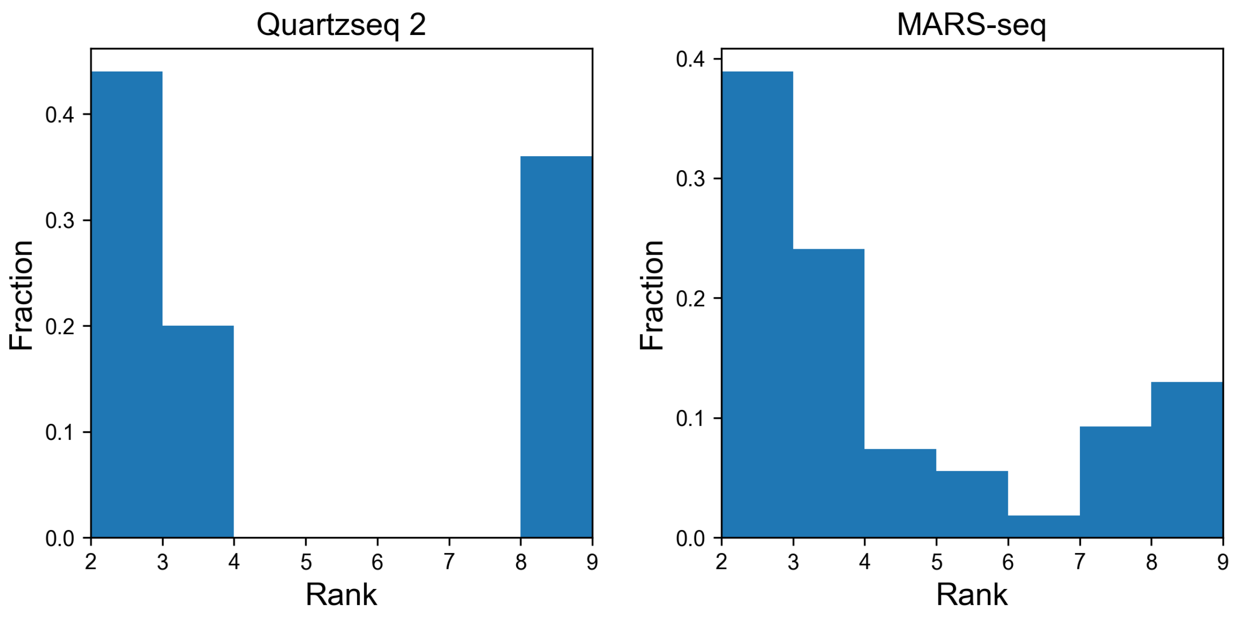


FIGURE S5: Distribution of second choices in label transfer. For each dataset in which we assessed accuracy of label transfer by SMB, we collect the rank of affinities of the correct class when an assignment could not be performed (*i.e.* the highest affinity was for the "Unknonwn" label)

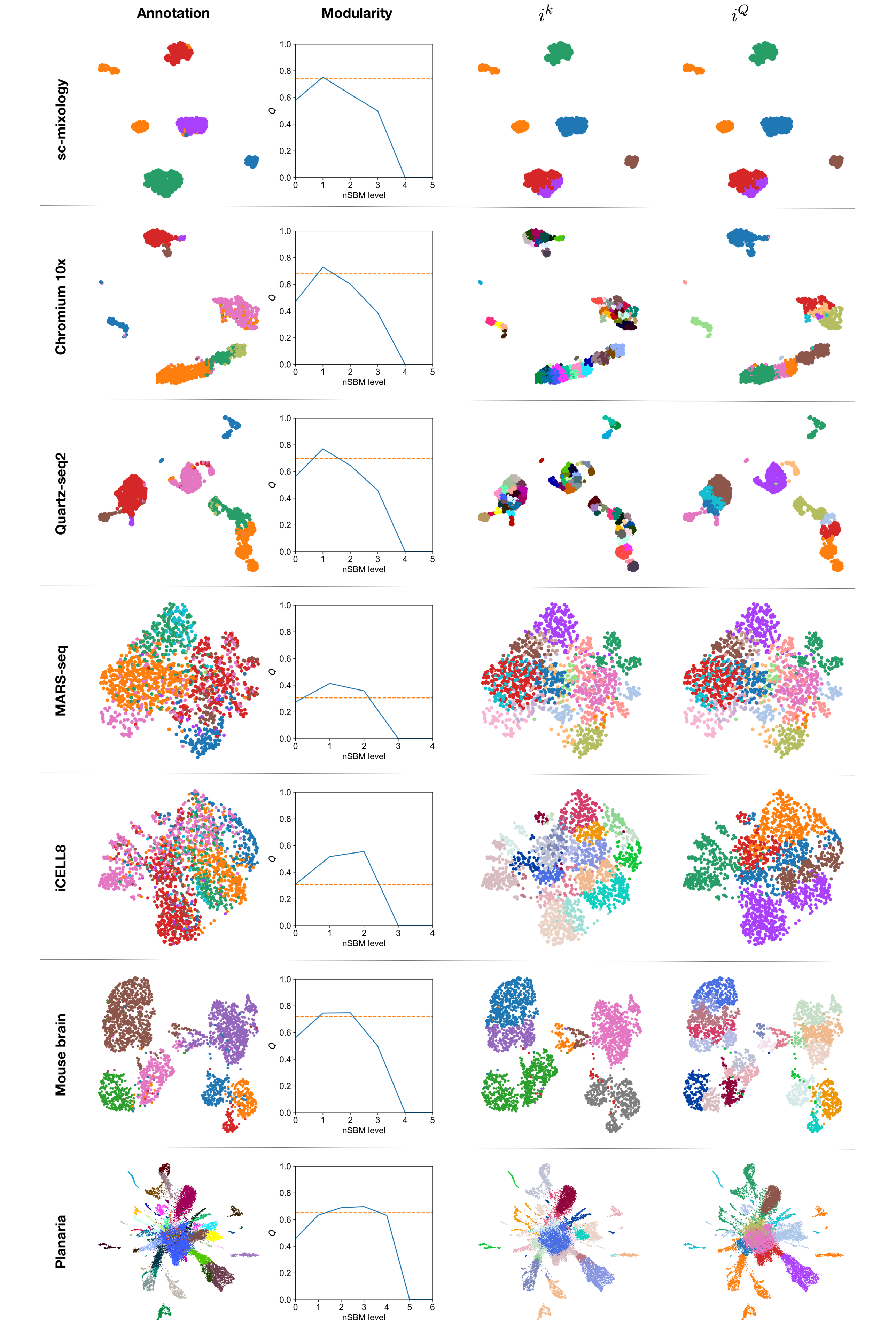


FIGURE S6: Choice of the optimal hierarchy level. Datasets analysed for choice of the most relevant hierarchy level are represented here, from top to bottom in the same order as in Table 2. For each dataset we report the UMAP embedding coloured by the annotation given in the corresponding manuscript, [illegible]
